# Nonhuman primates exhibit error type-dependent post-error slowing across modalities

**DOI:** 10.64898/2026.09.23.753764

**Authors:** Nami Kaneko, Mmesoma Nwokolo, Katherine Conen, Defne Buyukyazgan, Brent Stewart, Theresa M. Desrochers

## Abstract

Mistakes make up our daily lives. Once a mistake has been committed, it is common to pause before you continue. This slowing is called post-error slowing (PES). Although several theories have been proposed to account for PES in humans, findings across cognitive tasks show variation in how PES occurs, and it is unclear how this phenomenon may translate across modalities to a key model organism of human cognition, macaque monkeys. To determine whether monkeys exhibit PES, we tested three rhesus macaque monkeys in components of cognitive sequential decision-making tasks that used either eye tracking or touch screens to report responses. Here we show that monkeys exhibit non-adaptive, error type-dependent PES across modalities. Task structure and context also modulated PES in the eye tracking task, and no modality improved task performance following PES. These results suggest that monkey PES is a combination of cognitive and non-cognitive related responses to error that are modulated by task structures such as error type and task context. Overall, the flexibility in monkey PES is comparable to that of humans, further supporting the importance of monkeys as a translational model in investigating the mechanisms of PES.

## Introduction

Every day we make mistakes. After a mistake, it is common to pause when performing the next action. For instance, adding sugar into your stir-fry instead of salt may cause you to pause the next time you add an ingredient. This phenomenon is called post-error slowing (PES) and has been characterized as the slowing in reaction time on a correct choice following an error compared to the reaction time following a correct choice (Laming, 1979; P. M. Rabbitt, 1966). This post-error behavior is not reserved to humans, however, with limited evidence supporting that nonhuman primates (NHPs) also exhibit an increase in reaction time after an error (Phillips & Everling, 2014; Purcell & Kiani, 2016a). However, it is unknown whether this phenomenon is similarly flexible in NHPs as it is in humans. Specifically, whether PES is flexible and adaptive across modality and context, has yet to be studied in NHPs. Examining how PES manifests in NHPs is critical to investigating how behavioral mechanisms of post-error behavior may be conserved across primates.

PES has been found in humans in multiple response modalities but has only been found in eye tracking tasks in NHPs. Humans exhibit PES in studies that include eye tracking (Purcell & Kiani, 2016a), button pressing (Balogh & Czobor, 2016), and grasping movements (Ceccarini & Castiello, 2018), indicating PES exists similarly across multiple modalities. In a comparative eye tracking stop-signal task both humans and NHPs exhibited PES (Phillips & Everling, 2014). In NHPs, PES has been found in an eye tracking oculomotor switch task and stop-signal task with results suggesting that the mechanisms underlying post-error adjustments are similar in both humans and NHPs (Godlove et al., 2011; Purcell & Kiani, 2016a). While human PES reveals the flexible and multi-modality exhibition of PES, PES in NHPs has only been found in eye tracking modalities. Investigating if PES translates to other modalities, like it does in humans, provides insight into the flexibility of PES and similarities across primates.

PES, in both its presence and the effect on performance, is not a uniform phenomenon. PES has been found in a variety of cognitively complex tasks and reaction time tasks. Stroop tasks (Regev & Meiran, 2014), flanker tasks (Fischer et al., 2018; Houtman & Notebaert, 2013; Saunders & Jentzsch, 2012), stop signal tasks (Ide & Li, 2011), reaction time grasping tasks (Ceccarini & Castiello, 2018), visuomotor tasks (McDougle, 2022), and visual search tasks (Steinhauser et al., 2017), have all been found to exhibit PES. Task type alone does not determine whether PES is exhibited. For example, in flanker tasks PES has been found to increase performance (Fischer et al., 2018) as well as decrease performance (Houtman & Notebaert, 2013) and in some cases, PES fails to emerge completely (Saunders & Jentzsch, 2012). While changes across tasks are variable and unclear, changes within the same task also reveal the inconsistencies on the effect of task on PES (Ceccarini & Castiello, 2018). Taken together, differences in PES across and within various task types reveals that PES is not a single uniform phenomenon.

Variability across task types suggests that task type alone is insufficient to explain when and how PES emerges. Previous work suggests that PES is influenced by the interaction of task structure features with task type. Changes to the cognitive demand (Regev & Meiran, 2014; Schroder et al., 2013), the learning stage (McDougle, 2022), the overall performance (Fievez et al., 2022), whether the errors are conscious (Saunders & Jentzsch, 2012; Schiffler et al., 2017), the type of error (Eben et al., 2023), and the type of feedback (Ali et al., 2024) in a given task modulate PES. However, task-specific changes do not affect PES uniformly suggesting an interaction with task type. While the mechanisms in which these factors effect PES remain unclear, together they indicate that PES is flexible across tasks.

Investigating how PES is modulated within tasks in the current literature has proven difficult with conflicting evidence of how PES is modulated, but sequences can provide a unique window into understanding the effects of task structures. PES has been found in a discrete sequence production task (Ruitenberg et al., 2014) which finds that after an error the entire sequences of button presses have an overall slower RT, with the first press slower than the rest of the button presses in the given sequence. This task informs how PES affects sequence production using a motor task. Abstract sequences, however, have yet to be studied in the context of PES. Abstract sequences are sequences that are represented by a rule, such as AAAB or same-same-same-different. These sequences are not bound by time or stimuli, meaning an abstract sequence, such as brushing your teeth, can take 2 or 10 minutes and the type of toothbrush or toothpaste does not matter (Desrochers et al., 2022). As a novel way of investigating PES, abstract sequences provide a unique opportunity to observe PES in sequences dissociated from motor sequences. Since abstract sequences are governed by a single rule but contain sub-tasks, we can gain perspective in understanding how PES manifests under certain task features within the same paradigm, providing a clearer avenue for investigating the complexity of what modulates PES.

There are two main ways to consider what PES is: what is occurring during the prolonged reaction time post-error and what the effect of PES is on performance. First, many theories attempt to explain what occurs during PES. The *Cognitive Control Account* of PES is the most popular framework of PES (Botvinick et al., 2001; Dutilh et al., 2012), wherein an individual spends time assessing their mistake to increase their performance following an error (Gehring & Fencsik, 2001). Increasing response caution after an error implicates error awareness and a consideration of the error in a cognitive fashion (Botvinick et al., 2001; Danielmeier & Ullsperger, 2011). In contrast to the *Cognitive Control Account,* the *Orienting Account* operates on a non-cognitive framework. Under the *Orienting Account,* PES allows for “orienting” back to the task after a self-committed distracting error (Notebaert et al., 2009; Steinhauser et al., 2017).The *Orienting Account* proposes that PES is less of a cognitive control mechanism but rather a reorientation to the task after an “oddball” event, a surprising or expectation violating error that draws attention away from the task (Barcelo et al., 2006; Castellar et al., 2010; Notebaert et al., 2009; Van Der Borght et al., 2016). The *Cognitive Control Account* is the most popular explanation for PES with many cognitive tasks supporting this finding, however it is important to note that these theories may not be mutually exclusive, but instead modulated by task structures. Past work has suggested that PES may first be an orienting reaction that turns into a cognitive process after some time (Danielmeier & Ullsperger, 2011; Houtman & Notebaert, 2013; Jentzsch & Dudschig, 2009). The reality of PES appears to be a mixed phenomenon including both the *Cognitive Control Account* and the *Orienting Account* dependent on task structure.

Second, the *Functional account* and the *Non-functional account* (Houtman & Notebaert, 2013) describe the effects of PES on task performance. The *Functional accoun*t of PES describes PES as adaptive and positively affecting task performance and mastery. This account has been supported by primarily cognitive tasks that have observed a decrease in error rate following instances of PES (Ceccarini & Castiello, 2018; Dutilh et al., 2012; Saunders & Jentzsch, 2012; Steinhauser et al., 2017). The *Functional Account* does not account for all PES study results, however, with many studies finding that the *Non-functional Account* of PES best fits their results. PES in these cases either did not decrease error rate significantly (King et al., 2010) or contributed to an increase in error rate (ER) (Notebaert et al., 2009; P. Rabbitt & Rodgers, 1977; Van Der Borght et al., 2016). Together, the *Functional* and *Non-functional accounts* of PES describe the effect of error and subsequent delay on the performance of a task.

In the current literature, PES in an abstract sequence task in NHPs across modalities has yet to be studied, and the underlying reasoning for PES remains unclear. To examine whether NHPs fall into frameworks of PES that are more cognitively driven, or elicited more by surprise, this study investigates 1) how error types affect PES; 2) how task structure modulates PES; and 3) whether PES increases performance across error types and task structures. We examined PES in components of an abstract sequence choice task in three monkeys using eye tracking and touch screen modalities. We predicted that NHPs would exhibit PES across both modalities, similar to humans. Due to the mixed nature of PES, we predicted that task structure (error types, task context, and choice type) would affect PES differently. Lastly, we predicted that PES would increase task performance uniformly across error types, as previously seen with cognitive tasks. We found that PES exists across modalities in NHPs and is modulated by different task structures. We found error type to change the magnitude of PES across both eye tracking and touch screen tasks. Task context and choice position also affected PES but only at specific error types, indicating an interaction between the task structures. PES was not found to be adaptive and did not change the ER following PES across all modalities and structures. These results show that in components of an abstract sequence task, NHPs exhibit error type task structure dependent, non-adaptive PES. Variation in PES following these task changes suggest a complex system of processes occurring during PES during abstract sequences. While yet to be investigated in human abstract sequences, complex and flexible PES in NHPs reinforces similarities between humans and NHP post-error behaviors, suggesting a similarly robust behavioral response following errors.

## Methods

### Participants

We tested three rhesus macaques, 1 male and 2 females (ages 14, 9, and 5, respectively, weighing between 6.5kg and 11.7kg, during data collection). Following the principles of the 3R’s (Reduction, Refinement, Replacement) in animal research and standard practices in nonhuman primate neuroscience studies, the sample size was slightly larger than the minimum number of subjects as established in the field (two) to ensure that a minimum number learn the task and advance to future experiments. Though the overall number of participants was relatively small, robust statistical power was ensured through a large number sessions and trials per participant in the dataset. A total of 206 sessions were analyzed (157 Monkey W, 21 Monkey S, 28 Monkey V). With each monkey completing an average of hundreds of trials per day (875 Monkey W; 1,300 Monkey S; 680 Monkey V), the total dataset included approximately a total of over 180,000 trials (∼137,375 trials Monkey W; ∼27,300 Monkey S; ∼19,040 Monkey V). All effects were investigated in individual participants to examine consistency and ensure results were not driven by outliers.

The male, Monkey W, had a head holder implanted while under anesthesia during a sterile surgical procedure. All procedures, training, and data collection, followed the NIH Guide for Care and Use of Laboratory Animals and were approved by Institutional Animal Care and Use Committee (IACUC) at Brown University.

### Abstract Sequence Choice Task

The Abstract Sequence Choice Task used in this study is based on a previous no-report abstract sequence viewing task (Yusif Rodriguez et al., 2023). In the current task, subjects were tasked to identify what comes next in a 4-item abstract sequence by making a choice between two stimuli. Monkey W made choices using eye tracked saccades, while Monkey S and Monkey V indicated their choices using touch screen touches. While the overall goal of the task was the same, because monkeys S and V had learned components of what Monkey W had learned, we will describe the eye tracking and touch screen versions of the task separately.

#### Eye Tracking Abstract Sequence Choice Task

##### Stimuli

The eye tracking abstract sequence choice task was presented and recorded using MonkeyLogic (Hwang et al., 2019). Eye position was monitored using video eye tracking (Eyelink 1000, SR Research). Fractals were generated using MATLAB with custom code (Kim & Hikosaka, 2013; Miyashita et al., 1991). Across the sessions included in this study, each session included 6 distinct fractals that were reused for three sessions every 10 sessions. All stimuli were presented at the center of a 50.8 cm diagonal screen on a gray background (0.5 red, 0.5 green, 0.5 blue), with a fixation dot (0.1° circle) that was present on the screen during the task superimposed on the images and not present during the intertrial interval. To initiate fixation, the gaze was limited to 1.4 degrees of visual angle (DVA) until the trial began. This window was hidden from the subject and operated as a boundary to determine adequate fixation. Once fixation was acquired and the trial began displaying stimuli, the fixation boundary expanded to 1.7 DVA. When choice stimuli were presented, the fixation dot moved from the center to the two choice locations superimposed on the fractals. The fixation window for each choice was 1.5 DVA to ensure the selection of the choice was deliberate.

##### Trial Structure

Following the rule AAAB, Monkey W selected the next fractal in the sequence in two conditions: regular choice and forced choice trials (**Fig. 1A, B)**. Across both trial types, the fractal sequence of same-same-same-different or AAAB remained the same. Monkey W was tasked with making a decision at position 2 (A**A**) or position 4 (AAA**B**). The trial position type was pseudorandomized throughout the task ensuring equal numbers of position choices. Each trial consisted of one choice. Fixation was required to initiate the trial and then the first fractal appeared. Among regular choice position 2 trials (**Fig. 1A**), after the first fractal image presentation (300 ms) and an interstimulus interval (300 ms), two fractals appeared, indicating a choice. With the two fractals on the screen, the fixation dot remained in the center of the screen for a 500 ms choice delay when the monkey had to maintain fixation at the center. After that delay, the fixation dot disappeared from the center and two dots appeared at the center of each of the available choice fractals and indicated the go-cue. At this go-cue, Monkey W made a choice by making a saccade to and fixating (250 ms) on one of the presented fractal choices. Correct choice side was pseudorandomized throughout the task, ensuring equal numbers of choices on either side. The trial ended after the correct choice stayed on the screen for 250 ms as feedback for the monkey and fixation was initiated for the next trial.

**Fig. 1.**
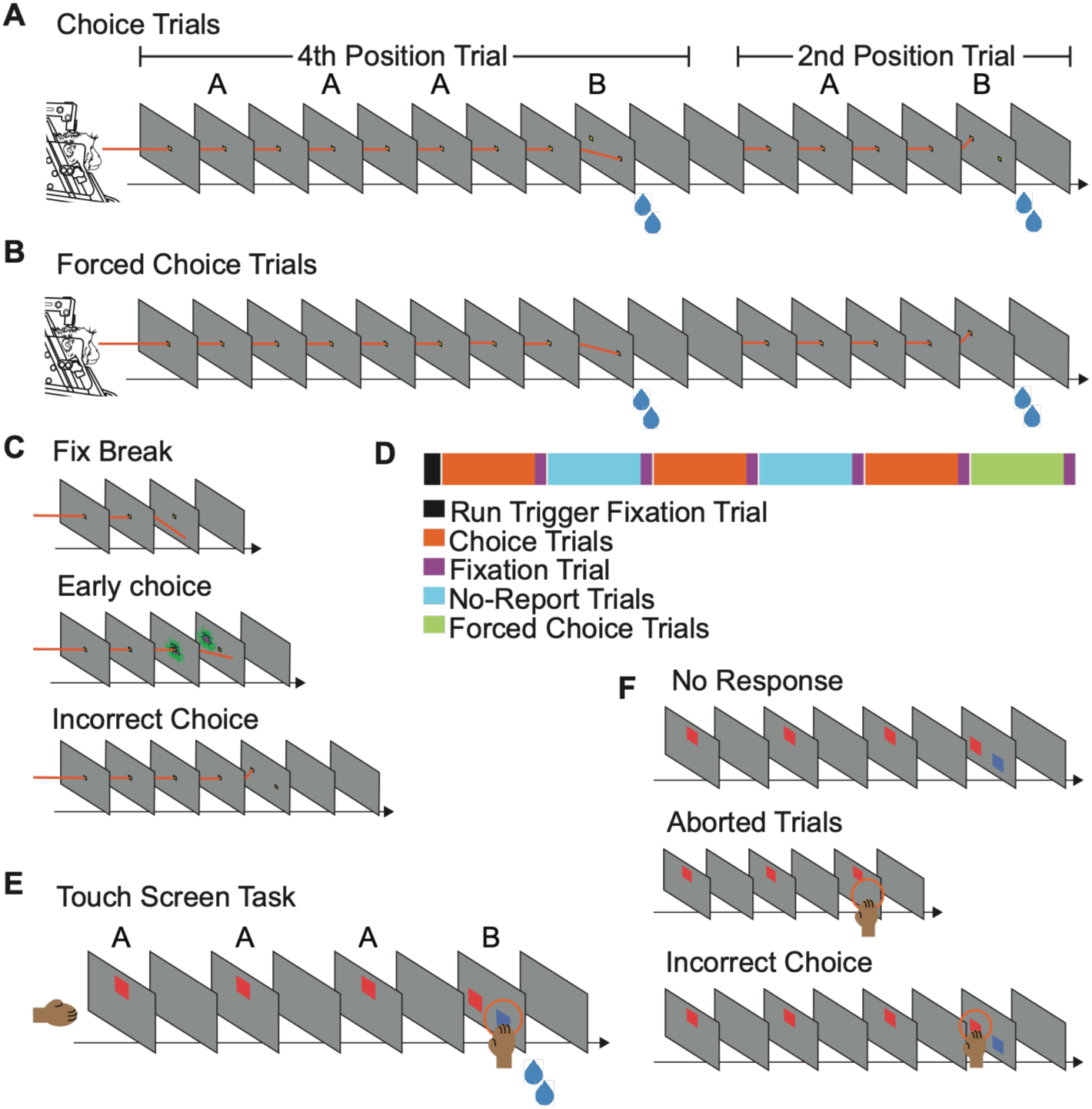
Abstract sequence choice task. **A.** Example trials of the eye tracking task (Monkey W) choice trials with choice position 4 (AAAB, left) and position 2 (AA, right) **B.** Example eye tracking task forced choice trials illustrating position 4 (AAAB, left) and position 2 (AA, right) trials that mirror the choice trials but only the correct choice is provided to the monkey. **C.** Illustration of the three possible types of errors in the eye-tracking task: fix break, early choice, and incorrect choice. **D.** Example run structure. Blocks of trials ended with a 14 s fixation-only trial (no fractals shown on screen). **E.** Example trial of the touch screen task (Monkey V and Monkey S). **F.** Illustration of the two possible error types in the touch screen task: incorrect choice and aborted trials.

Forced choice trials (**Fig. 1B**), had the same structure as regular choice trials except only the correct choice fractal was displayed on the screen instead of a choice between two fractals. With just the correct choice on the screen, Monkey W saccaded to the correct fractal and the overlaid fixation dot.

For a correct response of choosing “same” at position 2 trials or “different” on position 4 trials, the monkey was rewarded with two pulses of 60 ms of juice, after which the intertrial interval (ITI) began. ITIs were jittered in this task, ranging from 250 ms to 7,500 ms with a mean of 2,000 ms. There were three possible errors for choice trials in the eye movement task (**Fig. 1C**): fix break, early choice errors, and incorrect choice. In fix break errors, the monkey’s eye position left the allowable window around the fixation spot before the choice was seen, the trial terminated immediately, no juice was rewarded, and the ITI began. Early choice errors were when the monkey’s eye position left the allowable window around the fixation spot after the choices had been displayed but before the go-cue (during the 500 ms choice delay). After an early choice error, the trial was terminated immediately, no reward was delivered, there was a 2 s timeout with no fixation point on the screen, and then the ITI began. Lastly, if the monkey’s eye position left the fixation window after the choices were displayed at the appropriate time (i.e., after the choice delay), but the saccade was to the incorrect target, this error was deemed an incorrect choice. After an incorrect choice, no reward was delivered, the correct choice was displayed for 250 ms as feedback, there was a 2 s timeout, and then the ITI began. For forced choice trials, incorrect choices could not occur, therefore only fix break, and early choice errors were present during these trials.

Each run began with a fixation trigger block where after adequate fixation the experimenter would start the run. There were 7 blocks per run. Each block of the eye movement task included 28 trials with one 14-second fixation trial at the end of each block (**Fig. 1D**). There were three choice blocks in a run, amounting to 84 trials per run designated as choice trials. With one block of forced choice, 28 trials were designated as forced choice trials. The remaining 2 blocks were no-report AAAB sequence blocks. In the no-report blocks the monkey maintained fixation as the abstract sequence stimuli, also following the AAAB rule, appeared on the screen (Yusif Rodriguez et al., 2023). The monkey received two pulses of 40 ms of juice on 50% of the trials he successfully fixated (randomized). With a total of 6 blocks, 175 trials were included in each run. In a given session, Monkey W completed 5 runs on average.

#### Touch Screen Task

##### Stimuli

The touch screen task was similar to the eye tracking task (**Fig. 1E**). Monkey V and Monkey S used an in-chair tablet touch screen, the HomeBase system, created and developed by David Sheinberg and Ryan Miller at Brown University. Monkey V and Monkey S’s touch-screen version of the choice sequence task used simple colored squares as stimuli. For Monkey S, the colors were assigned at random following the rule AAAB. For Monkey V, there were five color pairs that used 10 colors, e.g., 1112, 3334, 5556, 7778, and 9990. Correct and incorrect choice colors were a different opacity to aid in training for both monkeys. Correct choices appeared at 100% opacity while incorrect choices ranged from 0% to 100% for Monkey V and 80% to 100% for Monkey S throughout the course of this dataset. Monkey S had two error types: aborted trials and incorrect choices. Touch windows, hidden to the subject, were the exact size of the colored square stimuli and were only used for Monkey S. If Monkey S touched anywhere on the screen while stimuli were being presented, outside of the choice time window, this action caused the trial to terminate, triggered a timeout of 2 s, and then proceeded to the ITI (1 s). Monkey V did not have aborted trials since she could not terminate the trial with out-of-bounds touches. Monkey V instead had no response trials where no choice was made throughout the entire trial. Monkey S and Monkey V did not have blocks or runs structured like Monkey W, instead each session was as many trials as they were willing to complete in a given day (680 trials on average per session for Monkey V and 1,300 trials per session for Monkey S).

##### Trial Structure

Following the rule AAAB, Monkey S and Monkey V completed trials for position 4 only and touched the screen to indicate their choice. Timings for each monkey were slightly different. For both, each trial began with the color square stimuli appearing on the screen with each color square appearing one at a time. For Monkey S, each stimulus was on screen for 180 ms with a 180 ms in between stimuli (inter-stimulus interval). For Monkey V, the stimulus on and off times fluctuated as training progressed, between 200 and 300 ms with an average of 200 ms. When choice position 4 was reached, the sample disappeared from the screen and two colored squares appeared as choices. Both monkeys touched the square on the screen to indicate their decision. Trials could be terminated by erroneous touches in Monkey S’s case or by no response for Monkey V. Feedback did not differ for correct versus incorrect choices; both choices disappeared from the screen and the 1 s ITI began.

### Data Analysis

Data analysis was conducted using MATLAB 2024a. Analysis used sessions where Monkey W, Monkey S, and Monkey V were attending to the task with consistent performance across sessions. For Monkey W, blocks where the subject was disengaged were omitted. The criteria for omitted blocks were seven or more sets of two consecutive trials with no fixation and six or more sets of three consecutive trials where the monkey broke fixation. These thresholds were empirically determined. Additionally, only sessions above 70% correct were included for all monkeys to ensure analysis only included trials where subjects were attending to the task. All trials were included in analysis for Monkey S. For Monkey V, blocks with bouts of longer than 10 subsequent aborted trials were omitted.

Reaction times (RT) for the eye tracking task were determined to start at the go cue (when the fixation dot moved to each possible choice fractal) and the touch screen RTs began at choice presentation. RTs for both modalities ended once the subject made their choice by saccading to or touching within the choice response circle. The correct choice trial where RT was measured was considered trial “N”. Trial N-1, was identified by the type of trial that came before N: correct, break fix, early choice, incorrect, abort, or no response. The combination of N-1 and N is a PES pair. N+1 is the trial that followed a PES pair and was used to examine the effect of PES on performance by determining the error rate (ER). Repeated measures anovas (RMANOVAs) were used to test the effect of error type, task type, and position type on RT and ER. Effects of PES were calculated by comparing the RTs of error-correct pairs to correct-correct pairs using t-tests pairwise comparisons. T-test comparisons of subtractions of error-correct RT by correct-correct RT were used to compare PES as well. Ratios (error-correct RT/correct-correct RT) were used to compare PES across individuals/tasks. P-values of comparisons were corrected for multiple comparisons using Bonferroni-Holm correction (Groppe, 2026). Analyses were carried out using custom scrips in MATLAB. RTs and ERs were plotted using notBoxPlot (Campbell, 2026). Overall ER was determined using all trials. We also calculated inverse efficiency scores (IES) for each monkey (RT ms/percent correct) to determine the speed accuracy tradeoff (Townsend & Ashby, 1978).

## Results

Three monkeys performed two versions of an abstract sequences task: one using eye tracking, and two using a touch screen (**Fig. 1A, E**). The eye tracking task required the monkey (Monkey W) to fixate during the presentation of stimuli until a choice appeared at position 2 (AA) or position 4 (AAAB) in an AAAB sequence. After a delay period, the go cue appeared allowing the monkey to saccade to a choice that fulfills the AAAB sequence. The touch screen task had Monkey S and Monkey V perform a part of the abstract sequence task used in the eye tracking version, the position 4 AAAB choice. A total of 206 sessions were analyzed (157 Monkey W, 21 Monkey S, 28 Monkey V). All monkeys performed well, above 70% correct in included runs. Error rates (ERs) were consistent for the trials used in analysis with an overall average of 15.5% for Monkey W, 23.24% for Monkey V, and 24.87% for Monkey S.

### NHPs exhibit PES across modalities

We first determined whether monkeys displayed PES across modalities. We measured the reaction time (RT) of a correct choice (N) that followed N-1, either a correct or error choice. Among all monkeys and both modalities, PES was exhibited. The RTs of error-correct pairs were longer than the RTs of the correct-correct pairs for the monkey performing the eye tracking task (Monkey W, **Fig. 2A**, t(156) = 8.3, p = 4.7 × 10⁻¹⁴, d = 0.66), and both monkeys performing the touch screen task (Monkey V, **Fig. 2B**, t(27) = 11, p = 9.8 × 10⁻^12^, d = 2.1; Monkey S, **Fig. 2C**, t(20) = 6.4, p = 3.4 × 10⁻^6^, d = 1.4).

**Fig. 2.**
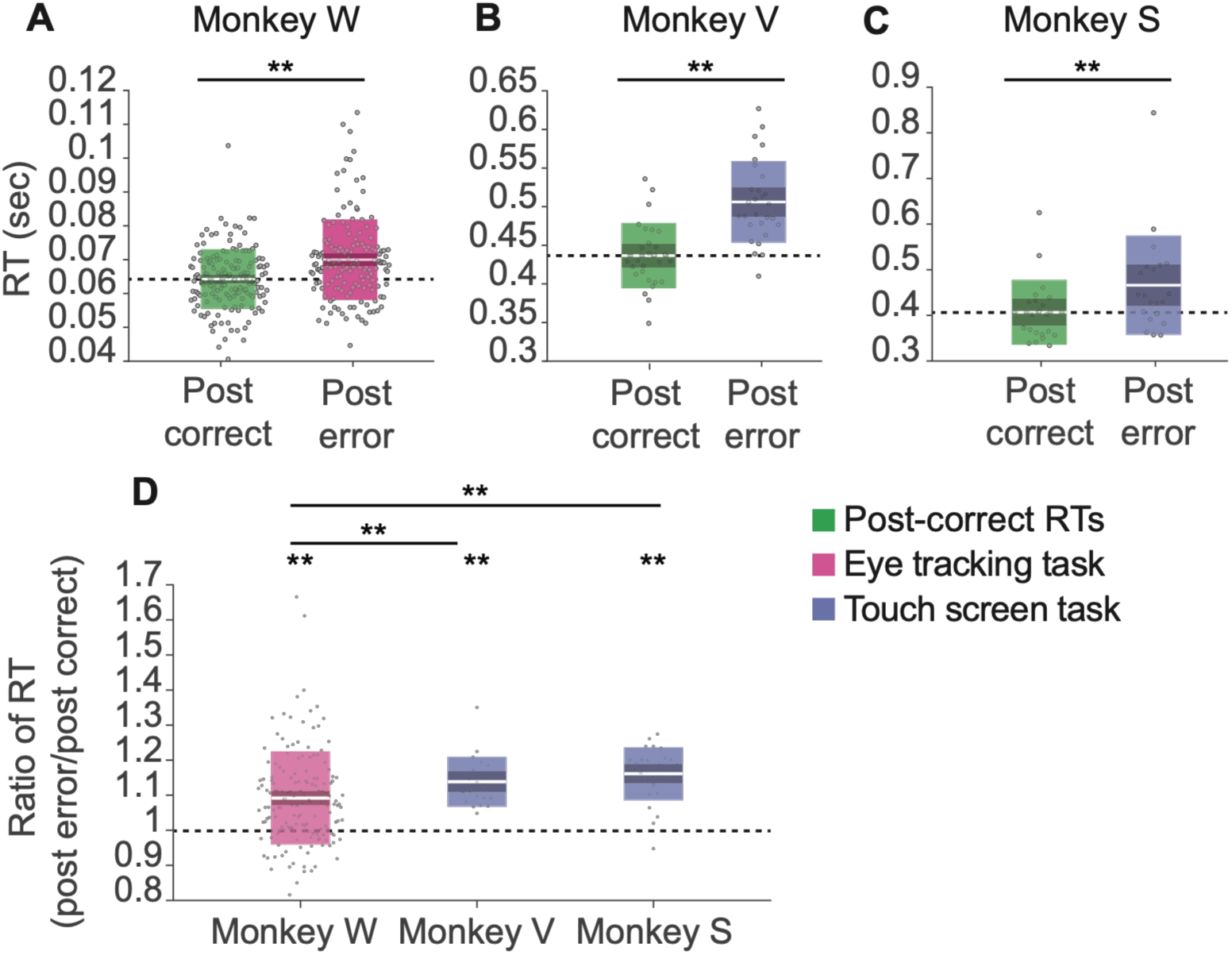
PES is exhibited in NHPs across modalities. **A**. RTs following correct choices (correct-correct) and following errors (error-correct). Each point is a median RT for that condition in a session. The dashed line indicates the mean of the correct-correct RT. The white line indicates the mean of each group, and the darker shade is the 95% CI. The box reaches 1 SD. n = 157 session medians. **B**. The same as A but for Monkey V, n = 28 session medians. **C.** Same as A but for Monkey S, n = 22 session medians. **D.** The ratio of error-correct/correct-correct of each monkey. P-values were adjusted for multiple comparisons using the Bonferroni-Holm correction. * = p < 0.5, ** = p < 0.001

To determine if the magnitude of PES was different between modalities and individuals, we calculated PES ratios (error-correct RTs divided by correct-correct RTs) for each subject and compared these ratios across subjects (**Fig. 2D**). On average, monkeys with the touch screen task exhibited similar PES ratios which were higher than the ratio of the eye tracking task. Monkey V’s PES ratio was significantly greater than Monkey W’s (t(182) = 180, p = 2.0 × 10⁻^207^, d = 13). Monkey S’s ratio was also significantly greater than Monkey W (t(175) = 180, p = 3.0 × 10^-203^, d = 14), but Monkey V and Monkey S were not significantly different from each other (t(46) = -1.1, p = 0.28, d = 0.16). Together these results indicate that PES was exhibited for both modalities and greater in the touch screen task than the eye tracking task.

### PES is modulated by task context

To investigate the effect of task structure on PES, we examined the differences in PES (trial N) across error types in the trial before (N-1) across both modalities. We also explored the effect of task type and choice position within the eye tracking task (which was not possible in the touch screen task). We measured the effects of each task structure by examining the RT of N and the ER of N+1.

#### PES is affected by error type

To investigate whether PES is modulated by the type of error, we determined whether errors preceding the correct choice (N-1) affected the RT of N.

##### Eye tracking Task

We separated the error-correct pair by the preceding error type. In the eye tracking task there were three error types: “fix break”, when the monkey’s eye position left the fixation window prior to the choices being displayed; early choice errors or “early”, when a choice was made too early during the choice delay period; and “incorrect”, when the wrong choice was selected. Across all sessions overall, fix break-correct pairs consisted of 10% of all pair types, early-correct pairs consisted of 5%, incorrect choices were 2% while the majority of pairs were correct-correct pairs at 83% (**Fig. 3A**).

**Fig. 3.**
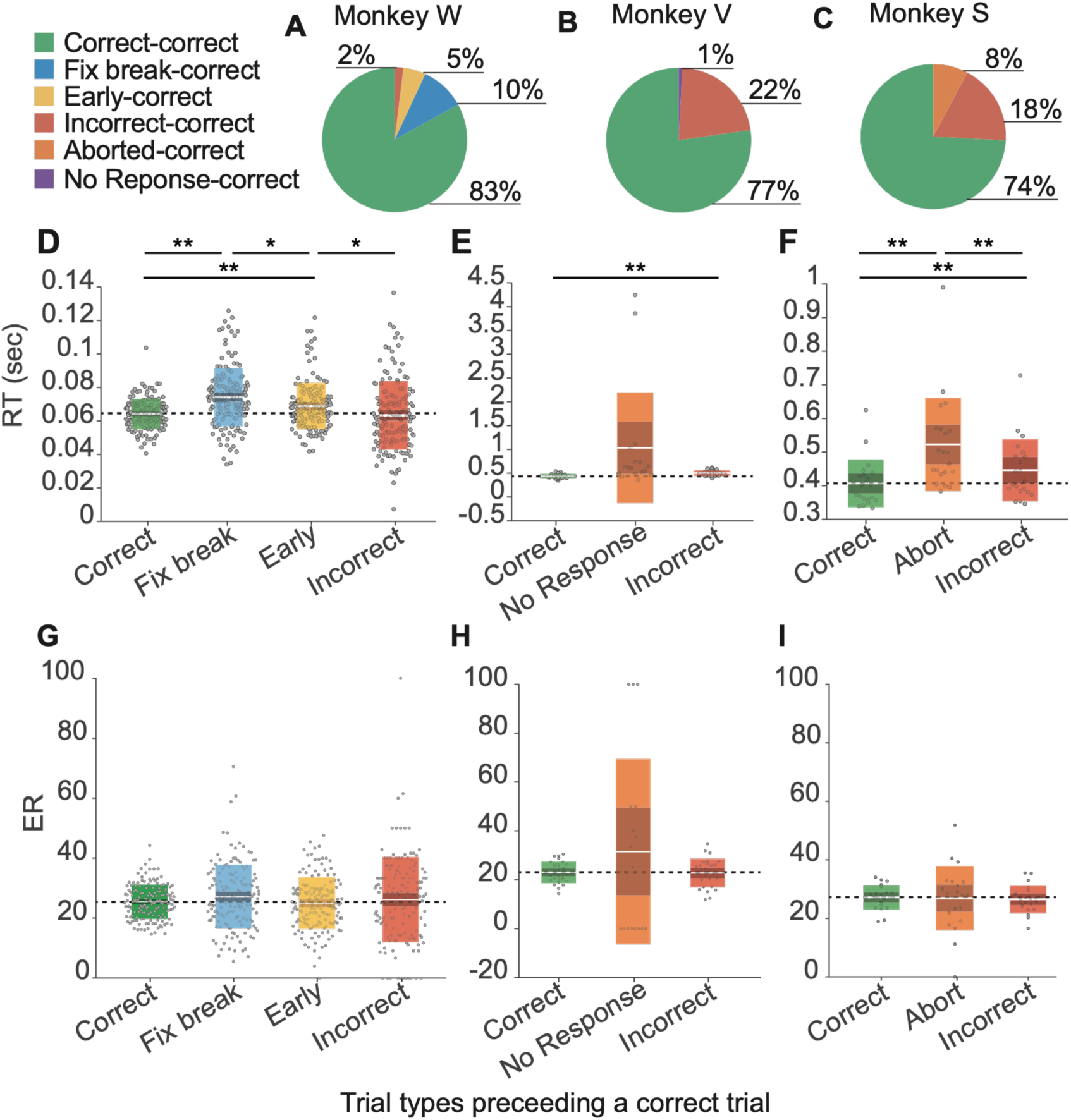
Error type affects PES RT and ER. **A.** Distribution of correct choices following different trial types in eye tracking task (Monkey W). Average percentages across sessions, rounded to the nearest whole number displayed on the pie chart. **B.** Same as A, but for touch screen task Monkey V. **C**. Same as A, but touch screen task Monkey S. **D.** RTs of N following trial types (N-1). The darker colored stripes at the center of the bars indicate 95% CI. The white line indicates the mean and the larger box is 1 SD from the mean. Monkey W, n = 157 session medians. **E.** Same as D but Monkey V, n = 28 session medians. **F.** Same as D but Monkey S, n = 22 session medians. **G.** ER of N+1 by trial type (N-1). Monkey W, n = 157 sessions. **H.** Same as G but Monkey V, n = 28 sessions. **I.** Same as G but Monkey S, n = 22 sessions. * = p < 0.5, ** = p < 0.005. P-values were adjusted for multiple comparisons using the Bonferroni-Holm correction.

Direct comparisons (**Table 1**, **Fig. 3D**) revealed that correct choices preceded by fix breaks and early choice errors exhibited longer RTs relative to correct-correct choices and therefore exhibited PES (p = 1.9 × 10^-15^, p = 1.7 × 10^-5^, respectively). PES was not found following incorrect choices (p = 0.52). To determine whether PES of each respective error type differed, we calculated the RT difference between the N trials on correct-correct pairs and the N trials for each error-correct type pair. Fix breaks had the largest magnitude of PES compared to early choice errors (t(156) = 3.9, p = 1.4 × 10^-4^, d = 3.1) and incorrect choices (t(156) = 6.5, p = 1.1 × 10^-9^, d = 0.52). Early choice errors had a larger magnitude of PES relative to incorrect choices (t(156) = 3.4, p = 8.6 × 10^-4^, d = 0.27).

**Table 1.** Results of Post-hoc T-tests of differences in RT by error type. P-values were adjusted for multiple comparisons using Bonferroni-Holm correction. RT comparisons are made using the correct choice (N) following the labelled trial type (N-1).

| Comparison | | $T(df)$ | $p$ | Cohen's $d$ |
| --- | --- | --- | --- | --- |
| **Eye tracking (Monkey W)** |  |  |  |  |
| Correct | Break fix | 9.1(156) | $1.9 \times 10^{-15}$ | 0.73 |
| " | Early | 4.8(156) | $1.7 \times 10^{-5}$ | 0.38 |
| " | Incorrect | -0.65(156) | 0.52 | 0.05 |
| Break fix | Early | 3.9(156) | $4.2 \times 10^{-4}$ | 0.31 |
| " | Incorrect | 6.5(156) | $5.5 \times 10^{-9}$ | 0.52 |
| Early | Incorrect | 3.4(156) | $1.7 \times 10^{-3}$ | 0.27 |
| **Touch screen (Monkey V)** |  |  |  |  |
| Correct | No response | 2.1(16) | 0.096 | 0.52 |
| " | Incorrect | 11(27) | $3.0 \times 10^{-11}$ | 2.1 |
| Incorrect | No response | -1.9(16) | 0.096 | 0.47 |
| **Touch screen (Monkey S)** |  |  |  |  |
| Correct | Abort | 6.1(20) | $1.4 \times 10^{-5}$ | 1.3 |
| “ | Incorrect | 6.6(20) | $6.0 \times 10^{-6}$ | 1.4 |
| Incorrect | Abort | -4.1(20) | $5.7 \times 10^{-4}$ | 0.89 |

##### Touch screen Task

To investigate whether error type modulated PES in the touch screen task, we analyzed the RT of the correct trial (N) following either a correct or error trial (N-1). The touch screen task included incorrect trials for both Monkey S and Monkey V. Monkey S also had aborted trials and Monkey V had no response trials. Incorrect trials were trials where the monkey touched the incorrect choice. Monkey S could abort trials by touching anywhere on the screen while Monkey V’s no response trials were caused by not attending to the task. These trial types are both considered “opted out” trials where the monkey did not make a choice, whether that be through touching the screen to end the trial or abstaining from the trial completely. Monkey V had 22.6% of incorrect choice pairs, 0.3% of no response trial pairs, and 77.1% correct pairs (**Fig. 3B**). Monkey S had 18% of incorrect choice pairs, 8% of aborted trial pairs, and 74% correct pairs (**Fig. 3C**).

We examined PES for each error type in each monkey performing the touch screen task. For Monkey V, incorrect-correct trial RTs were significantly greater than correct-correct trials (**Fig. 3E**, **Table 1**, p = 3.0 × 10^-11^), therefore exhibiting PES. No response-correct trials were not reliably different from correct-correct trials (p = 0.096). However, the magnitude of PES (correct-correct RTs subtracted from error-correct RTs) was not different between the two error types (t(16) = 1.9, p = 0.071, d = 047). Monkey S, who could abort the trial by touching anywhere on the screen outside the choice window, showed PES on both aborted trials and incorrect choices (**Fig. 3F**, **Table 1**, p = 1.4 × 10^-5^, p = 5.7 × 10^-4^, respectively). The magnitude of PES was greater for aborted trials compared to incorrect trials (t(20) = 4.1, p = 5.7 × 10^-4^, d = 0.89). Therefore, all monkeys across tasks showed differences in PES following different error types. While Monkey W did not exhibit PES following incorrect choices, Monkey V and Monkey S did. Monkey S also experienced PES following aborted trials. Overall, PES changed depending on the error type across tasks and individuals.

#### PES across error types did not increase performance accuracy

To continue investigating the effects of error type (N-1) on PES, we tested whether error type of PES affected task performance in support of the *Functional* or *Non-functional Account*. We compared the ER of the trial following a correct-correct pair, and the ERs following error-correct pairs (n+1). The *Functional Account* predicts an increase in performance following instances of PES while the *Non-functional account* predicts no change or a decrease in performance.

##### Eye tracking task

Results from the eye tracking task support the *Non-functional* account of PES across error types. For the eye tracking task, N-1 had no effect on N+1 ER (**Fig. 3G**, F (3, 468) = 1.6, p = 0.19, η2 = 0.01). Post-hoc tests comparing the effect of N-1 trial types showed no change in ER when compared to the ER following correct-correct pairs (**Table 2**). To determine whether the lack of change in ER was driven by a speed accuracy tradeoff of the overall performance rather than an effect of PES itself, we calculated an inverse efficiency score (IES) (Townsend & Ashby, 1978). Monkey W was not only fast but incredibly accurate and exhibited no speed accuracy tradeoff overall with an IES of 89 (avg RT = 73.6ms)/(1-avg ER = 0.155). While the range of IES is dependent wholly on the task itself, lower numbers indicate less speed accuracy tradeoff. The high speed and high accuracy performance show that the constant ER across N-1 error types is not dependent on any tradeoff. Together, these results support the *Non-functional Account* of PES.

**Table 2.** Results of Post-hoc T-tests of differences in ER by error type. *P*-values were adjusted for multiple comparisons using Bonferroni-Holm correction. ER comparisons are made using the correct choice (N) following the labelled trial type (N-1).

| Comparison |  | *T(df)* | *p* | Cohen's *d* |
| --- | --- | --- | --- | --- |
| **Eye tracking (Monkey W)** |  |  |  |  |
| Correct | Break fix | 2.2(156) | 0.17 | 0.18 |
| “ | Early | -0.75(156) | 1.4 | 0.06 |
| “ | Incorrect | 0.65(156) | 1.0 | 0.052 |
| Break fix | Early | 2.2(156) | 0.17 | 0.17 |
| “ | Incorrect | 0.68(156) | 1.4 | 0.055 |
| Early | Incorrect | -0.99(156) | 1.3 | 0.079 |
| **Touch screen (Monkey V)** |  |  |  |  |
| Correct | No response | 1.0(16) | 0.96 | 0.25 |
| “ | Incorrect | -0.26(27) | 0.8 | 0.05 |
| Incorrect | No response | -1.0(16) | 0.96 | 0.24 |
| **Touch screen (Monkey S)** |  |  |  |  |
| Correct | Abort | -0.14(20) | 1.8 | 0.03 |
| “ | Incorrect | -0.58(20) | 1.7 | 0.12 |
| Incorrect | Abort | -0.16(20) | 1.8 | 0.0034 |

##### Touch screen Task

As in the eye tracking task, results from the touch screen task in both monkeys support a *Non-functional Account* of PES. Monkey V exhibited no difference in N+1 ER across N-1 trial types (**Fig. 3H**, F (2, 32) = 1.0, p = 0.38, η2 = 0.059) and the same was found for Monkey S (**Fig. 3I**, F (2, 40) = 0.057, p = 0.94, η2 = 0.0028). Pairwise comparisons also yielded no differences in ER (**Table 2**). Additionally, Monkey V and Monkey S had IES of 691 and 383, respectively. Compared to Monkey W, Monkey V and Monkey S were considerably slower and less accurate overall, reflecting a small speed accuracy tradeoff as well. Since performance did not improve, the touch screen task supports the *Non-functional Account* of PES.

#### PES is modulated by task type in the eye tracking task

To explore whether task context yielded variations in PES, we compared the forced choice block to the regular choice block in the eye tracking task. In the forced choice task, all timings, stimuli, and choice positions were identical to the choice task. The only difference between the two tasks was at choice selection. In the forced choice task, only the correct fractal image was displayed, requiring Monkey W to simply saccade to the correct side (**Fig. 1B**). To investigate whether PES was different from regular choice in this context, we replicated the same analyses from the regular choice task and compared them to forced choice.

To contextualize the frequency of errors we calculated the distribution of PES pairs of each error type amongst forced choice trials. Incorrect choices are not possible in this condition since only correct choices are displayed. There were two error types in the forced choice task: fix breaks and early choice errors. Fix breaks constituted 7% of all PES pairs, 1% were early choice errors, and 92% were correct-correct PES pairs.

Across all forced choice error types, overall post-error effects were exhibited. Error-correct trials had lower RTs than correct-correct trials (**Fig. 4A** left, F (2, 242) = 5.8, p = 0.0035, η2 = 0.046). There were differences among the error types in PES when examining the types independently (**Table 3**). Post-error speeding was found following early choice errors (p = 0.032) and RTs of early-corrects were significantly faster than fix breaks (p = 0.029). Fix breaks showed no PES (p = 0.31).

**Fig. 4.**
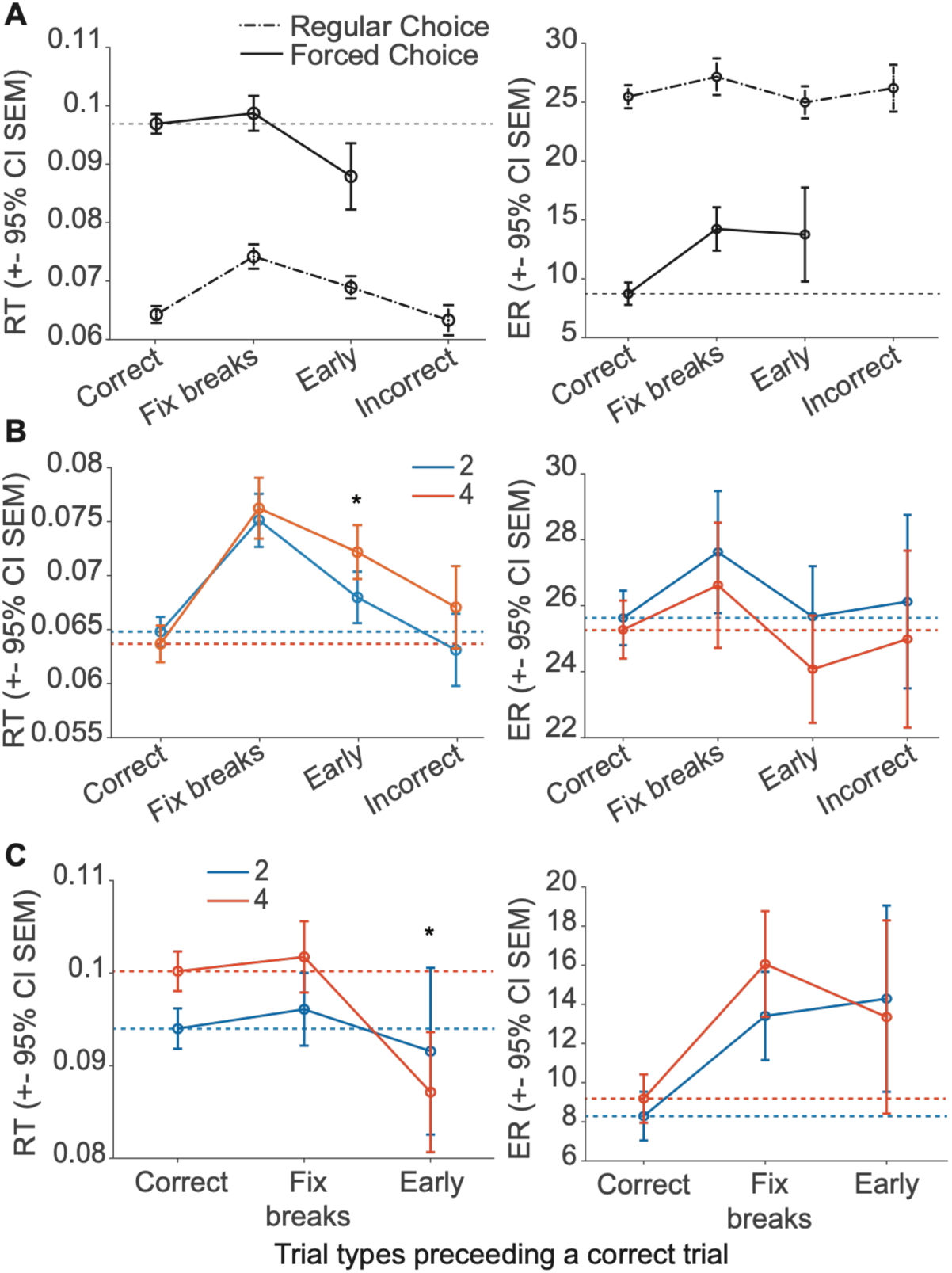
PES is modulated by task context. **A.** RT and ER of forced choice (solid line) compared to regular choice (dot-dashed line). **B.** RT (left) and ER (right) by choice position across PES pairs. Red lines indicate choice position 4 and blue indicates choice position 2. **C.** Forced choice by position by error. In all plots, horizontal dotted lines show mean RT (left) and ER (right) of correct-correct trials for visual comparison. n = 157, * = p < 0.5, ** = p < 0.005. P-values were adjusted for multiple comparisons using the Bonferroni-Holm correction.

**Table 3.** Results of Post-hoc T-tests of differences in RT by task context. Note. P-values were adjusted for multiple comparisons using Bonferroni-Holm correction. RT comparisons are made using the correct choice (N) following the labelled trial type (N-1).

| Comparison | | $T(df)$ | $p$ | Cohen's $d$ |
| --- | --- | --- | --- | --- |
| **Regular Choice** |  |  |  |  |
| Position 2 |  |  |  |  |
| Correct | Fix break | 7.9(156) | $2.8 \times 10^{-12}$ | 0.63 |
| “ | Early | 2.4(156) | 0.054 | 0.19 |
| “ | Incorrect | -0.88(156) | 0.38 | 0.071 |
| Fix break | Early | 3.8(156) | $7.2 \times 10^{-4}$ | 0.31 |
| “ | Incorrect | 5.2(154) | $3.1 \times 10^{-6}$ | 0.42 |
| Early | Incorrect | -2.3(154) | 0.054 | 0.19 |
| Position 4 |  |  |  |  |
| Correct | Fix break | 7.5(156) | $2.8 \times 10^{-11}$ | 0.60 |
| “ | Early | 6.0(156) | $5.5 \times 10^{-8}$ | 0.48 |
| “ | Incorrect | 1.5(153) | 0.14 | 0.12 |
| Fix break | Early | 2.1(156) | 0.099 | 0.17 |
| “ | Incorrect | 3.6(153) | 0.0019 | 0.29 |
| Early | Incorrect | -2.0(153) | 0.099 | 0.16 |
| **Forced Choice** |  |  |  |  |
| Correct | Fix break | 1.0(156) | 0.31 | 0.082 |
| “ | Early | -2.4(121) | 0.032 | 0.22 |
| Fix break | Early | 2.6(121) | 0.029 | 0.24 |
| Position 2 |  |  |  |  |
| Correct | Fix break | 0.90(154) | 1.1 | 0.07 |
| “ | Early | -0.057(96) | 1.1 | 0.054 |
| Fix break | Early | 0.64(96) | 1.1 | 0.065 |
| Position 4 |  |  |  |  |
| Correct | Fix break | 0.59(150) | 0.56 | 0.48 |
| “ | Early | -3.1(88) | 0.0075 | 0.33 |
| Fix break | Early | 1.6(85) | 0.24 | 0.17 |

To directly investigate if task context, forced or regular choice, affected post-error effects, we compared forced choice post-error effects from regular choice PES. Error-correct pairs of forced choice trials had significantly slower RTs on the N trial overall (F (1, 121) = 9.9, p = 0.0021, η2 = 0.075) and exhibited differing RTs across trial types compared to regular choice trials (F (3, 363) = 515, p = 2.1 × 10^-130^, η2 = 0.81). In summary, the forced choice context creates different post-error behavior from the regular choice context.

#### Task type affects performance following PES

Next, we examined ERs following the different PES pairs in forced choice to investigate whether the effect of post-error effects was also context dependent. Across all error types, error-correct trials had an overall increase in ER (**Fig. 4A** right, F (2, 240) = 4.6, p = 0.011, η2 = 0.037) and indeed, fix breaks-correct had significantly greater ERs compared to correct-correct ERs (t(156) = 5.6, p = 9.3 × 10⁻^8^, d = 0.45). However, this was not significantly different from early-correct ERs (t(120) = 0.80, p = 0.42, d = 0.073) which did not show any change in ER (t(120) = 1.8, p = 0.076, d = 0.16). Forced choice PES supports the *Non-functional Account* of PES.

To directly compare task context effects on post-error effects on ER, we compared the ER across trial types between forced choice and regular choice trials. We found that ER following any PES pair was much lower than regular choice trials (**Fig. 4A**, F (1, 120) = 266, p = 3.2 × 10^-32^, η2 = 0.69) and emerged differently by error type (F (3, 360) = 26, p = 4.9 × 10^-15^, η2 = 0.18). While significantly different in their ERs, these results support the *Non-functional Account* of PES across both task contexts.

#### PES is modulated by choice position in the eye tracking task

The eye tracking abstract sequence choice task included two types of choices, position 2 and position 4 (**Fig. 1A**) while the touch screen task only included a position 4 choice (**Fig. 1E**). At position 2, the correct choice was to choose the same stimulus as seen at the previous position (“same”, AA). In contrast, at position 4 the correct choice was to choose a different stimulus as was seen at the previous position (“different”, AAAB). While both kinds of choices were part of the same overall rule AAAB, they provide two different sub-task contexts within this paradigm. This situation was the case for both regular and forced choice trials.

To explore whether choice position in both regular choice and forced choice trials affected PES, we examined N RT and N+1 ER separated by choice position. Choice position in a PES pair was characterized as the choice position of N, the correct choice. N-1 could be any position. Each position type of N had similar proportions of error types, for both regular and forced choice trials. Regular choice PES pairs with N at position 4 constituted 47% of trials compared to 53% at position 2 over all sessions. N-1 for N at position 2 had 3% of trials as incorrect choices, 9% were early choice errors, 8% were fix breaks, and 80% were correct. N-1 trials for N at position 4 were 3% incorrect trials, 8% early choice errors, 7% fix breaks, and 82% correct choices. Forced choice PES pairs had 52% of N at position 2 and 48% at position 4. When N was at position 2 and position 4, the distribution was identical: 7% of N-1 were break fixes, 1% were early choice errors, and 92% were correct.

First for regular choice trials, we tested to see if the RTs of N changed depending on N’s choice position following different error types of N-1 (**Fig. 4B**). Direct comparisons revealed that position 2 showed PES following fix breaks (p = 2.8 × 10^-12^) and exhibited a marginal effect of PES following early choice errors (p = 0.054). Position 4 exhibited PES following fix breaks (p = 2.8 × 10^-11^) and early choice errors (p = 5.5 × 10^-8^). Within position 4, to see if PES following different errors were different from each other, we directly compared the RTs following fix breaks and early choice errors. RTs at position 4 following early choice errors were marginally different from fix break (p = 0.099) and incorrect choices (p = 0.099). To determine if PES of fix break and early choice error types were different by position we subtracted the RTs and found greater PES for position 4 choices following early choice errors compared to position 2 (t(156) = -2.9, p = 0.0044, d = 0.23). There was no difference between position for fix breaks (t(156) = -1.1, p = 0.26, d = 0.90). In summary, both choice positions exhibited PES following fix breaks and early choice errors, albeit position 4 had greater PES on early choice errors than position 2.

Similarly, to investigate whether forced choice PES was also affected by choice position, we directly compared the subtracted RTs of correct-correct from error-correct among position type 2 and 4 separated by error types (**Fig. 4C**). We found more post-error speeding for choice position 4 following early choice errors compared to choice position 2 (t(88) = 3.3, p = 0.0013, d = 0.35). Position 2 showed no post-error effects (p = 1.1). Among forced choice trials, position 4 showed the largest effect on post-error effects showing significantly distinct post-error speeding.

#### Choice position does not affect performance following PES

To explore whether choice position modulated the effect of PES or post-error speeding on ER, we measured the N+1 ER following Ns at position 2 or 4, separated by the N-1 trial type. For ERs of N+1 among regular choice trials, choice position had no effect (**Fig. 4B**, F (1, 151) = 1,6, p = 0.21, η2 = 0.010). N+1 ERs were also not different by error among the two N position types (F (3, 453) = 0.13, p = 0.94., η2 = 8.8 × 10⁻^4^). For forced choice we also saw no difference in N+1 ER by choice position (**Fig. 4C**, F (1, 59) = 0.0095, p = 0.76, η2 = 0.0016) and similarly, no effect of choice position by error type (F (1, 118) = 0.58, p = 0.56, η2 = 0.0097). These results again, support the *Non-functional Account* of PES.

## Discussion

The underlying cause and function of PES is still unclear with an emerging group of studies indicating a flexible and context-dependent form of PES (Danielmeier & Ullsperger, 2011; Jentzsch & Dudschig, 2009). To investigate how PES manifests in NHPs, we observed three rhesus macaques in a report-based abstract sequence task across eye-tracking and touch screen modalities. We found that NHPs exhibited PES across modalities, task structures influenced PES, and PES did not improve performance of the task. These results show the flexible nature of PES among NHPs and support the *Non-functional Account* of PES.

PES was found in NHPs across two modalities, eye tracking and touch screen, indicating that PES across modalities is conserved in NHPs. Humans show PES across eye tracking, button presses, and grasping movements (Balogh & Czobor, 2016; Ceccarini et al., 2019; Ceccarini & Castiello, 2018; Purcell & Kiani, 2016b). NHPs have only been shown to exhibit PES in eye tracking studies with results that show a similar capability of PES to humans (Phillips & Everling, 2014; Purcell & Kiani, 2016a). This study is the first to support that PES exists in touch screen for NHPs in addition to eye tracking. While the magnitude of PES differs across subjects, PES was observed across all modalities, illustrating its flexibility across species.

For all monkeys, different error types yielded differing effects on PES, supporting the claim that PES is mediated by error type. Prior work in humans has found that the type of error, error awareness, and performance monitoring contribute to PES, suggesting that PES by error type may be modulated by error type awareness (Chang et al., 2014; Damaso et al., 2020; Saunders & Jentzsch, 2012; Schiffler et al., 2017; Zhao & Cannon, 2026). This finding further supports that humans and NHPs react similarly to errors (Fu et al., 2023). However, whether awareness of errors contributes to PES is contested and it is suggested that PES does not improve task performance, concluding that error monitoring systems and PES are two distinct mechanisms (Dali et al., 2022). While the results of this study cannot say whether PES is affected by error awareness, different errors are indeed affecting PES.

PES was not associated with an improvement in performance, supporting the *Non-functional Account* of PES. To explain PES, the *Functional Account* and the *Non-functional Account* describe the effects of PES on task performance. The *Functional Account* which proposes that PES is an adaptive and functional mechanism predicts a decrease in ER and better performance following PES (Houtman & Notebaert, 2013). In the current study, performance was not improved across all subjects and modalities. Both the eye tracking and touch screen task showed no change in ER in the trial directly following PES regardless of error type, in line with other findings that support the *Non-functional Account* (King et al., 2010). Overwhelmingly, our results support a *Non-functional* account of PES.

PES in this study is not explained by the *Orienting Account*. In addition to the *Functional* and *Non-functional Account* framework, another way of investigating PES is examining what occurs during the slowing. Two such frameworks of what could be occurring during PES is described by the *Orienting Account* and *Cognitive Control Account*. The *Orienting Account* states that errors are considered “oddball” events, infrequent and surprising, requiring reorienting to the task after they occur (Barcelo et al., 2006; Castellar et al., 2010; Notebaert et al., 2009; Steinhauser et al., 2017; Van Der Borght et al., 2016). Overall, error-correct pairs are less frequent than correct-correct pairs for all subjects, which in theory follows the *Orienting Account* for infrequent errors causing PES. However, for the eye tracking task, the error with the largest number of trials caused the most PES. Furthermore, errors that occurred in the forced choice condition had no PES and in fact exhibited post-error speeding. Both the eye tracking and touch screen task exhibited PES for multiple error types but they were not as uncommon as “oddball” events as outlined by previous work (Notebaert et al., 2009). While we cannot say for certain some errors may be disrupting to the task and require orientation (perhaps for certain conditions and subjects), we can say that not all the PES exhibited in this study can solely be under the *Orienting Account* and PES may be attributed to something more cognitive.

PES in the eye tracking task is affected by task structures; namely the error type, the task context, and the type of choice. More specifically, we found that PES following early choice error trials was greatest at position 4 in the regular choice condition but caused post-error speeding in the forced choice condition. Choice position 2 versus 4 represent two different kinds of choices, cognitively and temporally, in one task. Additionally, the difference between regular choice and forced choice is whether Monkey W makes a choice between two options or saccades to the correct one. Therefore, our results may suggest a difference in cognitive context. Previous human work has found that PES is shaped by cognitive context (Fievez et al., 2022; Opdenaker et al., 2024; Regev & Meiran, 2014). Further, early choice errors and their effect on PES may indicate that inhibitory control plays a role in PES. Inhibitory control is necessary in making a decision. It is characterized by suppression and filtering of task-irrelevant information when recruiting decision-making mechanisms. Suggested to occur concurrently with orienting mechanisms (Wessel, 2018), deficits in inhibitory control diminish PES (Richard Ridderinkhof et al., 2011). Early choice error trials, which show the greatest post-error effects at position 4, are errors that occur before the delay period ends, possibly suggesting a lapse in inhibitory control. PES not only does not fully adhere to the *Orienting Account* but task structure specific modulations to PES suggest a cognitive quality to how PES manifests.

The present study is limited by sample size, and that the monkeys performing the touch screen task did not have choices at position 2. It will be important in the future to determine if the exploratory context-dependent effects observed in the eye movement task (i.e., the effect of position and forced choice) are also observed in different modalities. Given that other context effects (the type of error) similarly impact PES in both modalities, we hypothesize that similar effects will be observed. It will also be valuable to build on previous comparative work (Phillips & Everling, 2014; Purcell & Kiani, 2016a) and determine if humans performing the abstract sequence choice task show evidence of the conserved phenomenon of flexible PES.

Overall, these results show that NHPs and humans show similar dynamic post-error behavior. PES in an abstract sequence NHP task is exhibited across modalities, is task-context dependent, and is non-functional. PES was found to change depending on error type, task context, and position type suggesting that PES may be cognitive. This study is the first to identify PES in NHPs in an abstract sequence task across two modalities and suggests a paradigm of PES in NHPs that is modulated by task structure and context. Considering the similarities between humans and NHPs in this phenomenon, NHPs provide a valuable model for understanding PES and help uncover what neural processes underlie post-error behavior.

## Acknowledgements

Support: This work was supported by the National Institute of Mental Health (R01MH131615, T.M.D). Support was also provided by the National Science Foundation (BCS-2143656, T.M.D).

Thanks: We thank Matthew Maestri. Dr. David Sheinberg, Dr. Ryan Miller, Dr. M. Vanessa Rivera Núñez, Dr. Hannah Hyde, and Ellie McVeigh for their support and contributions to this work.

## Author Contributions

**Nami Kaneko:** Conceptualization, formal analysis, investigation, writing-original draft, writing-review & editing, visualization. **Mmesoma Nwokolo**: Conceptualization, formal analysis. **Katherine Conen**: software. **Defne Buyukyazgan:** investigation, software. **Brent Stewart**: investigation, software. **Theresa M. Desrochers:** supervision, project administration, funding acquisition, conceptualization, software, formal analysis, investigation, resources, writing-original draft, writing-review & editing

## Notes

### Competing Interest Statement

The authors have declared no competing interest.

